# Genome-wide strand asymmetry in massively parallel reporter activity favors genic strands

**DOI:** 10.1101/2020.08.26.269027

**Authors:** Brian S. Roberts, E. Christopher Partridge, Bryan A. Moyers, Vikram Agarwal, Kimberly M. Newberry, Beth K. Martin, Jay Shendure, Richard M. Myers, Gregory M. Cooper

## Abstract

Massively parallel reporter assays (MPRAs) are useful tools to discover and characterize regulatory elements in human genomes. Partly because enhancer function is assumed to be orientation independent with respect to each strand of the DNA helix, most reported MPRA results ignore stranded information. However, we find pervasive strand asymmetry of MPRA signals in datasets from multiple reporter configurations and in both published and newly reported data. These effects are reproducible across different cell types and in different treatments within a cell type, and are observed both within and outside of annotated regulatory elements. From elements in gene bodies, MPRA strand asymmetry favors the sense strand, suggesting that biological function related to endogenous transcription is driving the phenomenon. Similarly, within Alu mobile element insertions, we find that strand asymmetry favors the transcribed strand of the ancestral retrotransposon. The effect is consistent across the multiplicity of Alu elements in human genomes, and is more pronounced in younger, less diverged Alu elements. We find sequence features driving MPRA strand asymmetry and demonstrate its prediction from sequence alone. We see some evidence for both RNA stabilization and transcriptional activation mechanisms, and hypothesize that the effect is driven by natural selection favoring efficient transcription. Our results indicate that strand asymmetry, as a pervasive and reproducible feature, should be accounted for in analysis of MRPA data. More importantly, the fact that MPRA asymmetry favors naturally transcribed strands suggests that it stems from preserved biological functions that have a substantial, global impact on gene and genome evolution.

## Introduction

Spatiotemporal and quantitative control of transcript levels is a crucial aspect of essentially all biological processes in humans (Plank and Dean 2014; Schoenfelder and Fraser 2019). As such, finding the sequence elements that regulate transcription in human genomes and understanding the rules governing their effects are fundamental goals in human biology. For decades, these goals have driven a large amount of work, including both technology development (Johnson et al. 2007; Kwasnieski et al. 2012; Arnold et al. 2013; Gordon et al. 2020; Dekker et al. 2002; Patwardhan et al. 2009) and applications of those technologies to systematically find regulatory elements, including promoters, enhancers, silencers, and insulators (Ashe et al. 1997; Bell et al. 1999; Visel et al. 2009a; Dunham et al. 2012; Moore et al. 2020).

One key technological development has been the development of “massively parallel reporter assays” (MPRA), in which hundreds of thousands of elements are assayed in a single experiment for their ability to alter transcript levels. MPRAs take a variety of forms, but typically include the cloning of a diverse collection of short (∼200bp to ∼1.5 Kb) DNA elements into transcriptional reporter plasmid libraries (reviewed in Klein et al. 2019). These libraries are then transfected into cells that are subsequently subjected to sequencing.

One version of an MPRA, Self-Transcribing Active Regulatory Region sequencing (STARR-seq), places sequence elements within the 3’UTR portion of a gene in a plasmid construct that also includes a promoter (Arnold et al. 2013). The transcriptional enhancer effects of a given element, in a location that is downstream of the transcription start site (TSS) of the reporter DNA, can be directly quantified as each one contributes to its own abundance within the pool of plasmid-derived RNA. Another mode of MPRA, “survey of regulatory elements” (SuRE), involves placement of sequence elements in an upstream location relative to a gene in a promoter-free plasmid (van Arensbergen et al. 2019). These elements are linked to barcodes within the transcribed reporter, and their effects are quantified by measuring the abundance of their transcribed barcodes.

MPRAs of various types, including STARR-seq and SuRE, intrinsically allow for stranded information to be gathered; that is, the assignment of transcribed reporter fragment abundance to the tested plasmid DNA regulatory element can be oriented such that the direction of the DNA fragment can be tied directly to the orientation of the transcribed strand. However study designs based on fragment synthesis often ignore this capacity (i.e., if directional cloning is used and only one orientation is designed). In other experiments that use randomly fragmented DNA and non-directional cloning, stranded information is present within the resulting data but often largely ignored or assumed to have a small effect or an effect at only a limited number of loci (Ramaker et al. 2020; Muerdter et al. 2015; Sun et al. 2019; Schöne et al. 2018; Liu et al. 2017; Barakat et al. 2018; Wang et al. 2018).

The lack of emphasis on strand-specific effects is likely, at least in part, a consequence of the fact that enhancers, the target of many MPRAs, have traditionally been thought to be orientation independent. In fact, some propose that the ability to act in both forward and reverse orientations is a defining characteristic of enhancers (Andersson et al. 2014). Many individual enhancers have been shown to be approximately equally effective in both orientations in heterologous reporter assays (Visel et al. 2009b; Andersson et al. 2014; Dao et al. 2017; Mikhaylichenko et al. 2018; Klein et al. 2019). Further, many DNA-binding motifs for transcription factors (TFs) that bind to enhancers are also intrinsically symmetric (Sherwood et al. 2014).

However, the case for strand asymmetry of DNA function within MPRAs is, in general, strong. Indeed, gene transcription, perhaps the most fundamental of all biological functions encoded in DNA, is inherently stranded. Promoter activity is often directional, and even in cases where the activity is bidirectional, there is generally a bias toward one strand (Duttke et al. 2015; Andersson et al. 2015; Almada et al. 2013). Other properties, such as mutational correction, have also been shown to be strand-biased (Green et al. 2003). Furthermore, MPRAs may have features within their designs that predispose to strand asymmetry from non-regulatory effects. For example, In STARR-seq, the tested regulatory element is itself transcribed, implying that any sequence elements with strand-specific effects on RNA stability will lead to strand asymmetry in the data.

Here we show that strand asymmetry is a global feature of MPRA data. These effects are sequence-specific, and are highly consistent across cell-types and from independently sampled repetitive elements within a given experiment. Statistical modeling shows that the effects are predictable, at least in part, solely from primary sequence features. We find that MPRA strand asymmetry tends to favor the sense strand of genes encoded in human genomes and the transcribed strand of Alu insertions. These results are important for better design and interpretation of MPRA data. More importantly, the existence of a clear genic correlation suggests that the strand-specific signals being detected by MPRA reflect evolutionarily-guided mechanisms with widespread effects on genome sequences.

## Results

### Strand asymmetry is pervasive in MPRA signal

We considered four MPRA datasets in our analyses. We generated a STARR-seq (Arnold et al. 2013) library with a super-core promoter (SCP) (Addgene 71509) from sonicated BACs (bacterial artificial chromosomes) spanning a ∼1.2 Mb locus around the *HTT* gene (Supplementary Table 1). We assayed this library in four cell types: A549, BE(2)-C, HepG2, and K562 (Methods). We also generated a STARR-seq library using the promoter-less STARR-seq vector (Addgene 99296) from BACs in the *SORT1* gene locus in HepG2 cells (Methods).

In addition to these experiments conducted in our labs, we also obtained data from the Sequence Read Archive for a STARR-seq experiment using a fragmented whole genome library in A549 cells treated with dexamethasone for various durations (Johnson et al. 2018). Lastly, we obtained SuRE data from a reporter with the test element upstream of a promoter-less transcription start site (TSS) using libraries from four fragmented whole genomes in HepG2 and K562 cells (van Arensbergen et al. 2019).

For the three STARR-seq experiments, we aligned and processed the FASTQ files retaining both the alignment position and the strand orientation using a uniform pipeline (Figure 1, Methods). Due to the complexity of associating test elements to barcodes in the van Arensbergen et al. data from FASTQ files, we instead obtained stranded signal values from the bigWig files provided by the authors (GSE128325). All datasets were then mapped to a common set of 290 bp bins spanning the autosomes (Methods). To avoid ambiguity from terms such as “plus” or “minus” and avoid confounding genic strands with reference strands, we defined signal derived from reads matching the reference strand as “Reference” and those aligning to the reference reverse complement as “Complement”. From these signals, we calculated the value of Reference minus Complement (RMC) as a measure of strand asymmetry (Figure 1A). Because the van Arensbergen et al. data was processed differently, we found that division of RMC in this dataset by the maximum signal of either strand (RMC over Max) resulted in decreased variance across binned values within test categories (gene bodies, for example) and used this metric for all subsequent analyses on this dataset (see Methods).

**Figure 1.**
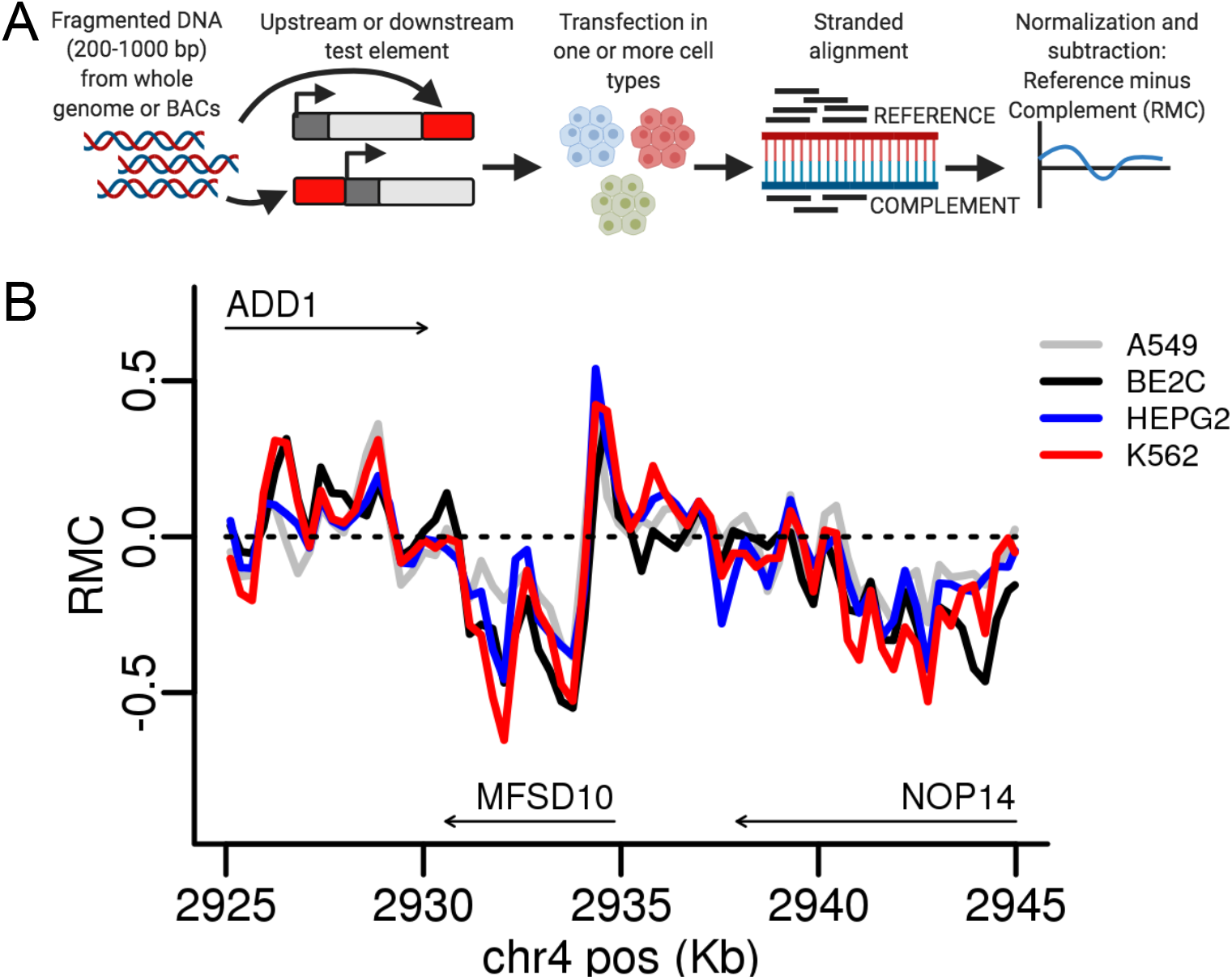
Derivation and example of RMC data. A. DNA test elements from either BACs or whole genome fragmentation are cloned downstream or upstream of the TSS in different experiments. After transfection in one or more cell types, reporter-derived RNA and DNA is harvested and sequenced. The reads are mapped retaining strandedness and the normalized signal Reference strand signal minus the Complement strand signal (RMC) is calculated. (Created with BioRender.com). B. RMC data from four cell lines is shown on a 20 Kb portion of chromosome 4 (hg38 coordinates). The arrows mark gene bodies and the direction of stable transcription.

We noticed that RMC from BAC-derived data in the *HTT* locus was consistent across the four cell types (Figure 1B). We explored whether this effect was pervasive across this dataset and the others, including those spanning the whole genome, and the BAC-derived library targeting the *SORT1* locus. For each experiment group (treatment, cell type, replicate) within each dataset, we calculated the reporter signal from the two strands separately. Within a dataset, we then calculated correlation between all strand-experiment group pairs (Methods). Finally, we clustered stranded reporter signal correlations for each dataset (Figure 2). The strand of the signal segregated with the first cluster clade, before even cell type (Figure 2A,C), technical replicates (Figure 2A, Supplementary Figure 1), dexamethasone treatment duration (Figure 2B), or genome donor (Figure 2C). We ruled out that biases in input reporter construct pools could be responsible for the strand asymmetry by comparing the sequenced DNA counts assigned to the two strands, finding high correlation (Supplementary Figure 2). Thus, strand asymmetry is the predominant feature driving global patterns of similarity of signals from MRPA datasets, being more prominent than any other technical (e.g., replicates) or biological (e.g., cell type) factor.

**Figure 2.**
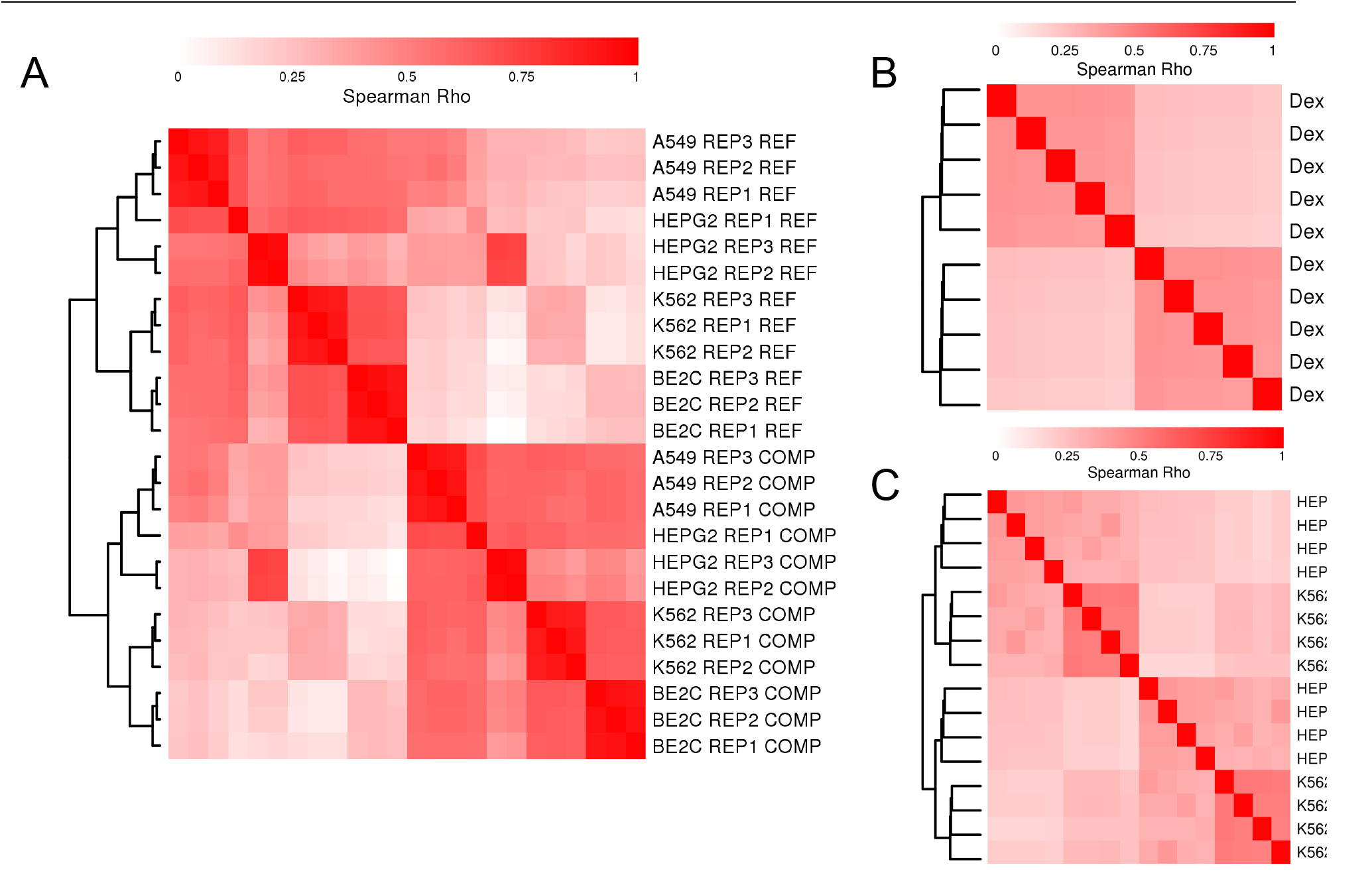
Clustering of MPRA signal by strand. Hierarchical clustering of Spearman correlation coefficients is shown for MPRA signal from A.) a pooled BAC-derived STARR-seq library spanning *HTT* gene in four cell types, B.) a whole-genome derived STARR-seq library in A549 cells from Johnson et al., and C.) whole-genome libraries from four donors in a promoter-less reporter system with an upstream test element in two cell types from van Arensbergen et al. All comparisons were made from binned data (see Methods).

In the Johnson et al. data, the dexamethasone treatment is proposed to activate regulatory regions, namely glucocorticoid response elements. We observed the same strand-driven clustering behavior both within and excluding likely regulatory regions defined by activating histone marks (Supplementary Figure 3 A,B). While the van Arensbergen et al. reporter aims to find promoter activity, we similarly observed strand-driven clustering in this dataset both within and excluding promoter regions, defined as −2000 to +500 bp from annotated TSS (Supplementary Figure 4). We were not able to find any genome segmentation, based on features like histone marks, promoters, or gene bodies, which removed the strand-driven clustering in either of the whole genome datasets. In fact, a randomly selected set of 1 million bins shows the same clustering pattern in both datasets (Supplementary Figures 3E, 4C).

### MPRA strand asymmetry correlates with gene bodies

Given the pervasiveness of the MPRA strand asymmetry, we sought to compare it to other genomic features displaying strand-specific effects. As the most obvious stranded genomic feature, we first considered gene bodies (defined by GTEx v8, including non-coding transcripts). Although the MPRA signal continues to cluster by strand both within and excluding gene bodies (Supplementary Figure 3 C,D), we hypothesized that RMC values would tend to favor a gene’s transcribed strand within gene bodies. For example, RMC should be positive in Reference genes (transcript sense to reference) and negative in Complement genes. To evaluate this, we divided autosomes into Reference gene bodies, Complement gene bodies, intergenic regions, or regions where transcripts from both strands overlap (Methods). Using linear regression, we found significant enrichment for positive RMC values in Reference gene bodies and negative values in Complement gene bodies compared to intergenic regions (all p < 2.2E-16, Table 1, Supplementary Figure 5 A-C). We found no significant difference between regions with transcripts on both strands and intergenic regions.

**Table 1.**
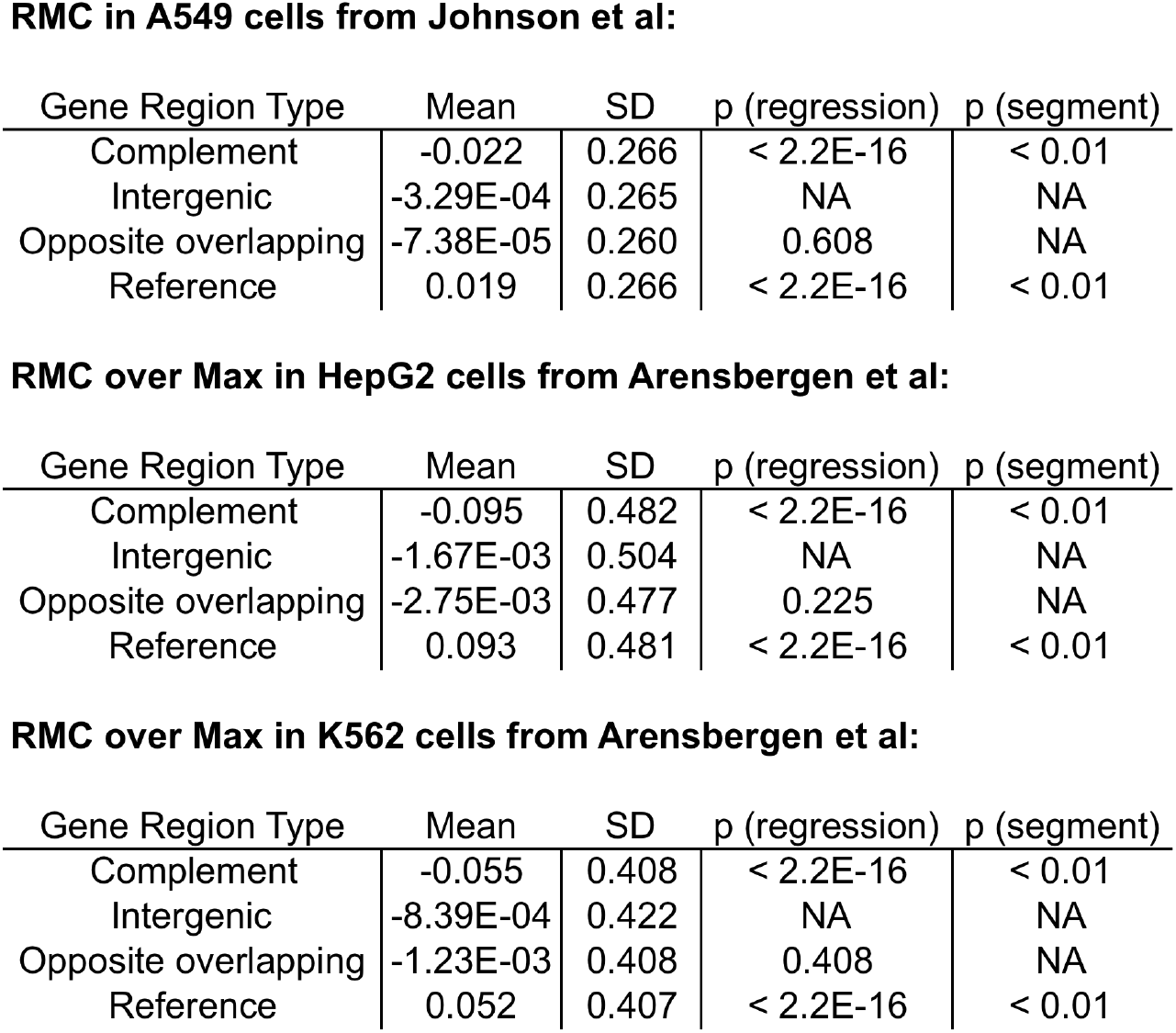
MPRA strand asymmetry shows significant and concordant association with gene body types. For these analyses the autosome was divided into Reference gene bodies, Complement gene bodies, regions with annotated transcription from both strands (Opposite overlapping), or intergenic (see Methods). The means and standard deviations for three datasets are presented. The p (regression) p-value is the linear regression p-value compared to Intergenic. The p (segmentation) p-value is the estimate of the significance of similarity of the HMMSeg segmentation from the dataset compared to the gene body type based on conditional entropies (see Methods).

**RMC in A549 cells from Johnson et al:**
| Gene Region Type | Mean | SD | p (regression) | p (segment) |
| --- | --- | --- | --- | --- |
| Complement | -0.022 | 0.266 | < 2.2E-16 | < 0.01 |
| Intergenic | -3.29E-04 | 0.265 | NA | NA |
| Opposite overlapping | -7.38E-05 | 0.260 | 0.608 | NA |
| Reference | 0.019 | 0.266 | < 2.2E-16 | < 0.01 |

**RMC over Max in HepG2 cells from Arensbergen et al:**
| Gene Region Type | Mean | SD | p (regression) | p (segment) |
| --- | --- | --- | --- | --- |
| Complement | -0.095 | 0.482 | < 2.2E-16 | < 0.01 |
| Intergenic | -1.67E-03 | 0.504 | NA | NA |
| Opposite overlapping | -2.75E-03 | 0.477 | 0.225 | NA |
| Reference | 0.093 | 0.481 | < 2.2E-16 | < 0.01 |

**Table 1.** MPRA strand asymmetry shows significant and concordant association with gene body types.
| Gene Region Type | Mean | SD | p (regression) | p (segment) |
| --- | --- | --- | --- | --- |
| Complement | -0.055 | 0.408 | < 2.2E-16 | < 0.01 |
| Intergenic | -8.39E-04 | 0.422 | NA | NA |
| Opposite overlapping | -1.23E-03 | 0.408 | 0.408 | NA |
| Reference | 0.052 | 0.407 | < 2.2E-16 | < 0.01 |

To further explore the association of MPRA strand asymmetry with gene bodies, we tested whether a segmentation of the genome by RMC (or RMC over Max) would be consistent with gene bodies. We used the HMMSeg tool (Day et al. 2007) segmenting into high (Reference-like) and low (Complement-like) states, at a range of transition probabilities, to all of the considered datasets (Methods). The resulting segmentations appear visually consistent with gene bodies across multiple datasets and cell types in the *HTT* locus (Figure 3, Supplementary Figure 6 A-C), and in the *SORT1* locus (Supplementary Figure 6D).

**Figure 3.**
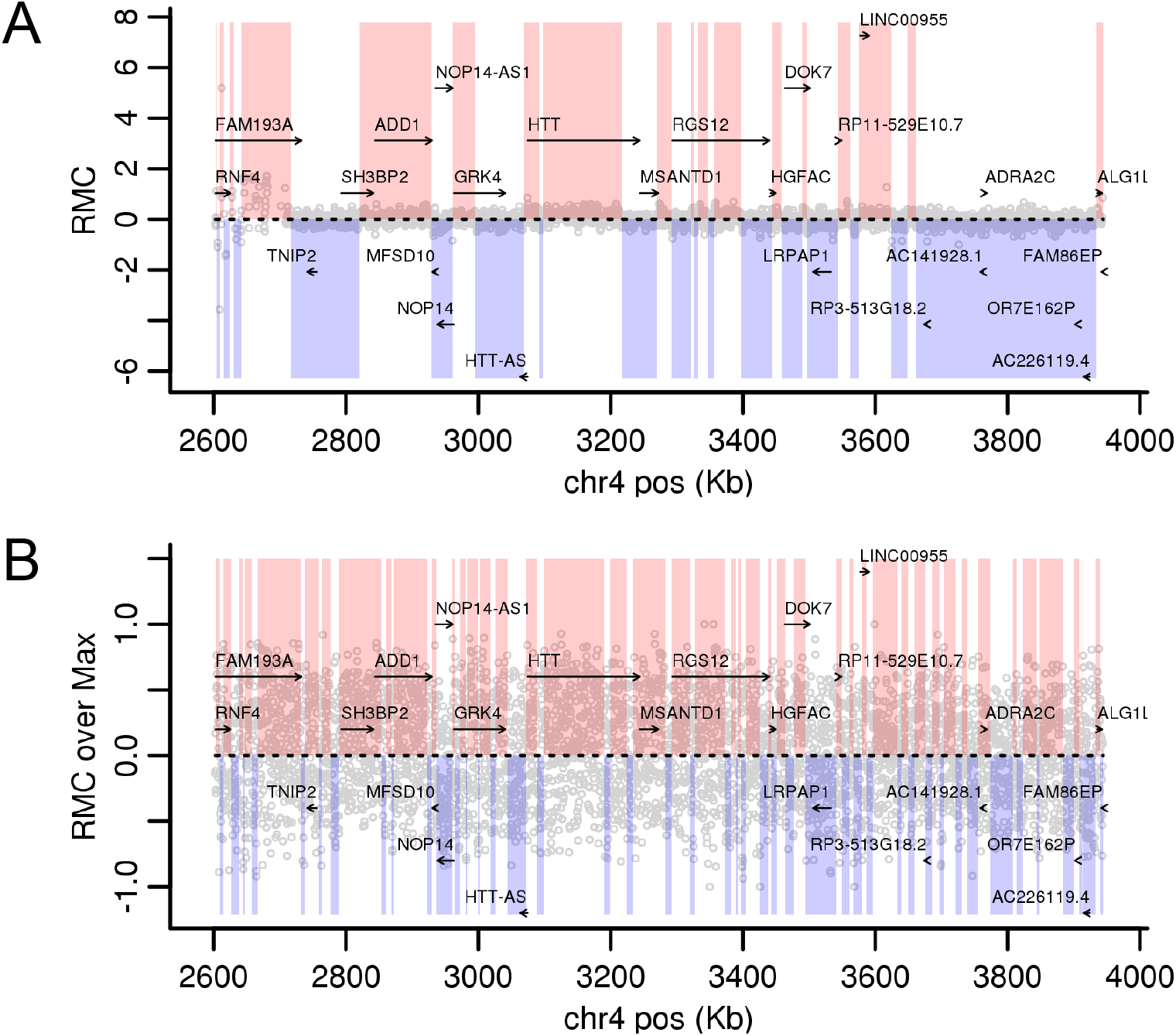
MPRA strand asymmetry is consistent with gene bodies. Values are plotted in the *HTT* locus for A.) RMC for STARR-seq signal from a library generated from BACs spanning the *HTT* locus in K562 cells, or B.) RMC over Max from Van Arensbergen et al. whole genome also in K562 cells. Gray dots are RMC bin values. Pink and blue blocks were assigned to Reference and Complement, respectively, using HMMSeg on each data set with a transition probability of 0.3 (see Methods).

To robustly evaluate the agreement with gene bodies genome-wide, we compared the HMMSeg derived autosome segmentations from Johnson et al. and van Arensbergen et al. datasets to gene bodies using a conditional entropy approach (Haiminen et al. 2007) (Methods). All datasets produced segmentations more similar to gene bodies than 1,000 randomly shuffled segmentations across a range of transition probabilities (Supplementary Figure 7, Table 1). While the segmentations did not match gene bodies perfectly, they classified a high fraction of Reference and Complement gene bodies correctly (Supplementary Figure 5 D-F). Intergenic and regions with transcripts on both strands were nearly evenly split between Reference and Complement segmentations (Supplementary Figure 5 D-F). In effect, genome-wide MPRA strand asymmetry data are able to accurately predict which strand is genic, including across intronic regions.

### Independent mobile element insertions are consistently strand-biased

Another strand-oriented genomic feature is repetitive regions derived from retrotransposons, which move through a transcriptional intermediate as part of their replication cycle. The most abundant such element in human genomes is the Alu element, present at more than 1 million copies in the reference human genome (Deininger 2011). We first examined the distribution of RMC values within Reference and Complement strand oriented Alus, and found a clear correlation (Figure 4), reminiscent of the genic strand bias (i.e., Reference transcribed Alu strand tends to have positive RMC).

**Figure 4.**
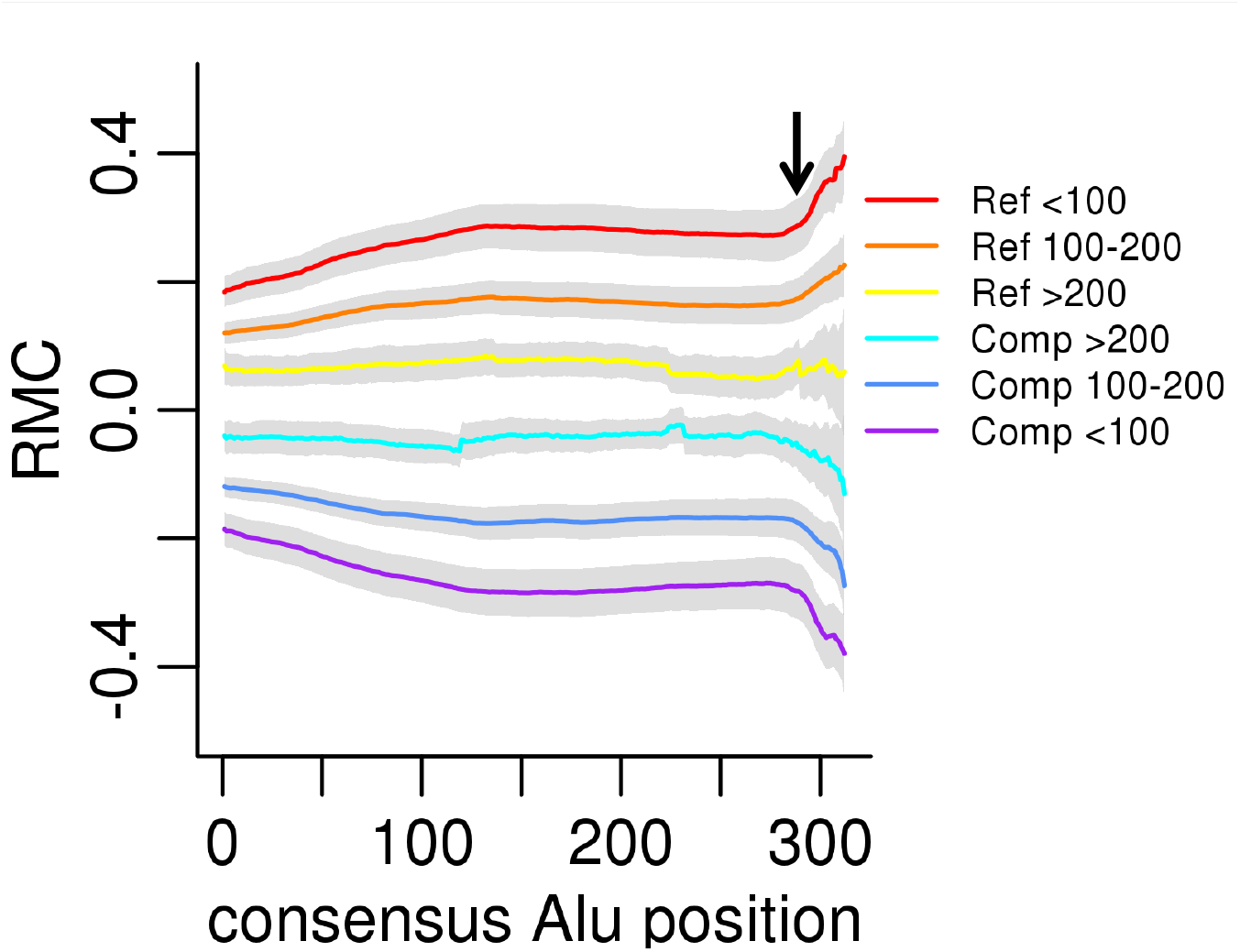
Alu divergence level effects strand asymmetry. From Johnson et al. data, the genome-wide median RMC (y-axis) for each annotated Alu consensus position (x-axis) is plotted for Reference- (Ref) or Complement- (Comp) oriented Alu insertions, grouped by levels of divergence (indicated in respective colors) measured by milliDiv units (e.g., < 100 corresponds to <10% divergence from the ancestral consensus, see Methods). The gray bands represent two standard deviations from the median. The black arrow indicates the start of the A-tail sequence.

We furthermore exploited the fact that Alus provide, in a sense, “biological replicates” of one another, being independently sampled genomic fragments that share similar sequence content. Within an annotated Alu, the genomic position can be mapped to a position within the Alu ancestral consensus sequence. We mapped each annotated Alu genomic base pair in data from Johnson et al. to positions in the Alu consensus sequence as determined by RepeatMasker (Jurka et al. 1996).

The resulting distributions of RMC values as a function of Alu consensus position indicate a clear pattern across the length of Alu sequences, with opposite strand orientation Alu-consensus positions mirroring one another. The 3’ end of Alu insertions are comprised of an A-tail, a feature that we see tends to intensify the degree of RMC in favor of the transcribed strand. Additionally, Alus are silenced after insertion and their sequence diverges from the ancestral sequence as mutations accumulate and are fixed over evolutionary time. We binned the Alu-RMC data by Alu divergence level and observed more intense effects from younger, less diverged Alu sequence (Figure 4). This observation is consistent with the sequence-specific nature of RMC within Alus. Younger, less divergent Alus are more similar to one another, and thus show more consistent effects, in contrast with older, more divergent and dissimilar Alus.

### Genomic sequence drivers of MPRA strand asymmetry

In order to find sequence features that might be predictive of strand asymmetry, we evaluated the correlation of monomer, dimer, and octamer frequencies within autosomal bins to RMC values from Johnson et al. We selected the significantly correlated k-mers and constructed a linear model trained on data from chromosome 1 (see Methods). This model accurately predicts RMC data on chromosome 2 (Figure 5A, p< 2.2E-16). We also used a sequence based predictive model of transcriptional activity using convolutional neural networks, Xpresso (Agarwal and Shendure 2020), to the *HTT* locus. We chose this smaller locus due to computational practicality. Xpresso predicted values significantly correlate to RMC data in that locus in four cell types (R^2^ = 0.11 to 0.46, Supplementary Figure 9).

**Figure 5.**
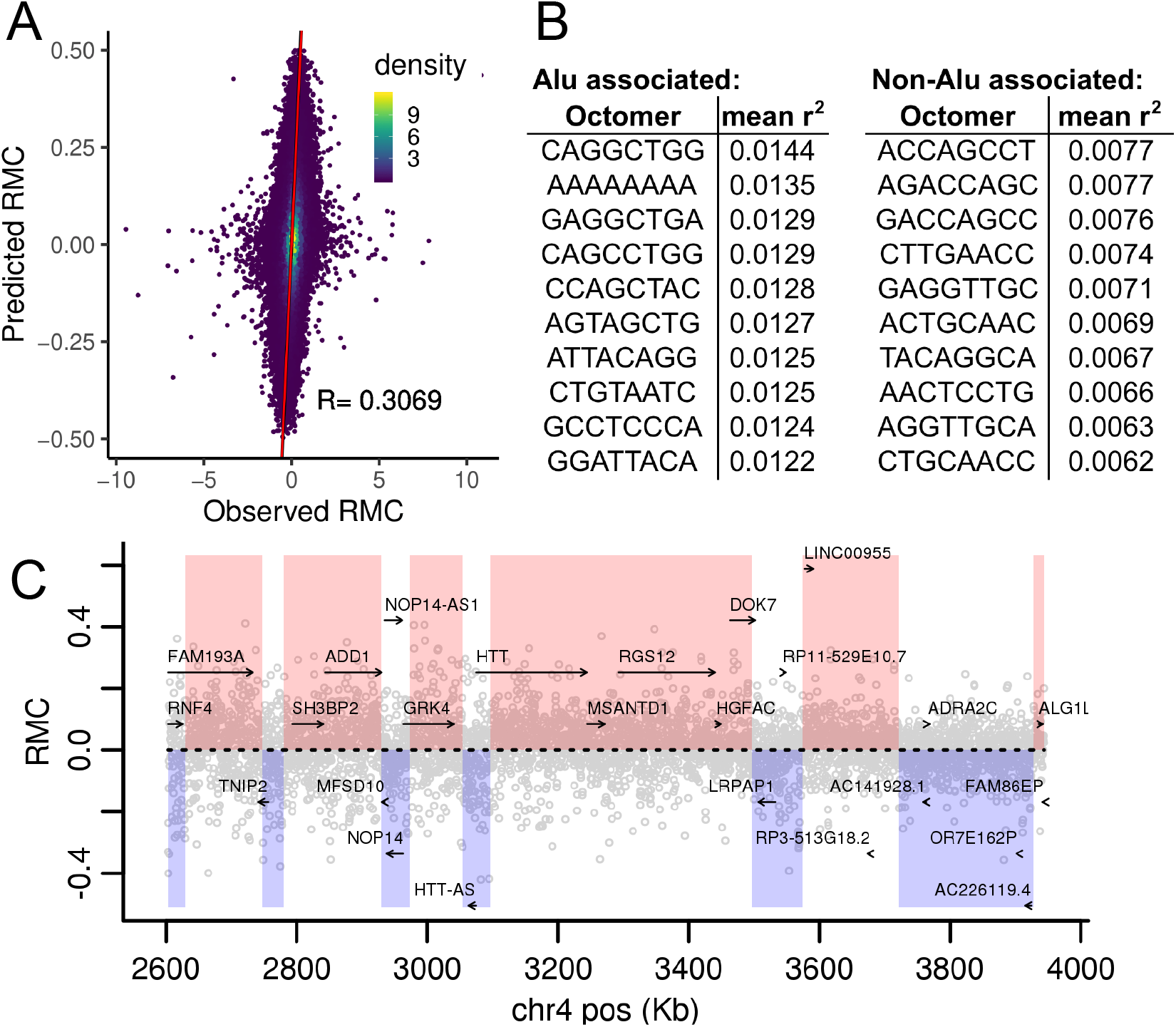
RMC predicted from sequence only. A.) A linear model based on monomer, dimer, and octamer frequencies was trained on chromosome 1 RMC data from Johnson et al. Model predicted RMC values for chromosome 2 are plotted versus actual values with Pearson’s R indicated. Red line represents the model fit. B.) The top ten most predictive octamers present in the AluY consensus sequence or those not are given. The r-squared is the mean of the sequence and its reverse complement. C.) Predicted RMC values in the *HTT* locus are plotted with gene bodies and HMMSeg derived segments. Gray dots are bin predicted RMC values. Pink and blue blocks were assigned to Reference and Complement, respectively.

Consistent with the strand asymmetry in RMC, a given k-mer’s reverse complement should have an equal but opposite effect. As expected, in linear models trained on only a single k-mer’s frequencies, k-mer performance closely matched that of its reverse complement, in opposite directions (Supplementary Figure 8). No palindromes yielded significant association. The top 54 most associated k-mer/reverse complement pairs all exist within the AluY consensus sequence. However many non-Alu k-mer pairs also strongly associated (Figure 5B, Supplementary Data 1). Only octamers appeared in the top ten k-mers for each category, although some monomers and dimers did yield significant associations (Supplementary Data 1).

Despite Alu-associated k-mer pairs driving some of the linear model’s accuracy, predicted RMC values are consistent with gene bodies in the *HTT* locus (Figure 5C). Also, HMMSeg segmentations of chromosome 2 derived from predicted RMC are more similar to gene bodies than any of 1,000 randomly shuffled segmentations across a range of transition probabilities (Supplementary Figure 10).

We annotated significantly predictive octamers by comparison to RNA-binding protein (RBP) and transcription factor (TF) motifs using FIMO (Grant et al. 2011). We found significant matches to RBP and TF motifs for some highly RMC-predictive octamers (Supplementary Table 2). However, matches were not found for most significantly predictive octamers.

## Discussion

In MPRA data generated from both fragmented whole genome DNA or targeted BAC pools, in STARR-seq vectors with and without the super-core promoter (SCP), and in SuRE vectors with an upstream, non-transcribed test element, we see pervasive and highly reproducible strand asymmetry in reporter signal. The effect persists over several cell types, from multiple donor genomes, and in differing drug treatments. Strand asymmetry is the predominant driver of clustering in all these datasets (Figure 2, Supplementary Figures 1,3,4).

The presence of strand asymmetry might be considered as merely an artifact of MPRAs were it not for its correlation to well-established stranded genomic features. Test elements derived from Reference-sense-strand genes have significantly positive RMC values while the converse is true for Complement-sense-strand genes in both datasets derived from whole genomic DNA (Table 1). Regions where stable transcription products are annotated on both strands yield RMC values near 0, perhaps indicating equilibrium between competing forces. Although MPRA strand asymmetry is pervasive, it appears to be organized coherently in gene bodies, allowing for segmentations via HMMSeg that are significantly similar to gene bodies (Figure 3, Supplementary Figures 6 and 7).

Test elements derived from Alu sequence show pronounced MPRA strand asymmetry (Figure 4). Ancestral Alu sequence has active retrotransposon activity that depends on transcription and interaction with LINE produced proteins (Deininger 2011; Mills et al. 2007). As Alus tend to degenerate after insertion and accumulate fixed changes over evolutionary time, they provide a natural mutagenesis experiment by allowing simultaneous assessment of ∼1 million independently inserted and evolved sequence fragments. We find that RMC values are clearly non-randomly distributed with respect to the transcribed Alu strand. Further, younger Alu elements, which are more similar to the consensus, display a greater magnitude of RMC at every Alu consensus position (Figure 4). We speculate that because Alus are compact features (∼300 bp) entirely transcribed and harboring regulatory elements, they represent “concentrated” units of MPRA strand asymmetric sequence.

It is expected that the data from van Arensbergen et al. displays strand asymmetry. The authors intended to detect promoter activity in their experiments and promoters are usually more active in one direction (van Arensbergen et al. 2019). We also note that because we used these files instead of processing FASTQs through the common pipeline used in all other presented datasets and used a modified strand asymmetry measure (RMC over Max), the van Arensbergen et al. data may be less comparable to the other datasets. Nevertheless, the van Arensbergen et al. data shows significant correlation with gene bodies similar to the Johnson et al. STARR-seq data despite the test element not being transcribed (Table 1). Also, segmentations by HMMSeg of the van Arensbergen et al. data are significantly similar to gene bodies, exhibiting the same regional coherence seen in Johnson et al. STARR-seq data (Figure 3, Supplementary Figures 6B, 7D-E). We speculate that in an upstream reporter configuration, gene body fragments increase the recruitment of RNA polymerase or other transcription initiators in a manner matching their genomic orientation. Previous research has highlighted the similarities between enhancer activity, enhancer transcription, and promoter activity, and it is possible that the effects we describe here are related to these phenomena (Mikhaylichenko et al. 2018). The bigWig files provided by the authors did not have sufficient resolution for an analysis of the Alu effect similar to the one performed on the Johnson et al. data.

Overall, we have shown that fragments of genomic sequence (∼200 bp to ∼1.1 Kb in the presented datasets) from gene bodies or Alus retain their strand asymmetry in an artificial reporter construct away from their native context of broader genomic organization, chromatin structure, nuclear localization, and three-dimensional conformation. This strongly suggests that the strand asymmetry is driven by sequence, as no other information is carried through to the reporters. We looked for nucleotide monomers, dimers, and octamers associated with strand asymmetry. Octamers contained within Alu sequence dominate the highest predictive k-mers, reflecting not only the potency of Alus’ effect but also their abundance in the human genome (Supplementary Data 1). We find some octamers significantly similar to RBP motifs and others to TF motifs (Supplementary Table 2). However, we found no “smoking gun” sequence class able to explain a large portion of the effect. Applying Xpresso, a tool that accurately predicts transcription level from genomic sequence only (Agarwal and Shendure 2020), yields predictions significantly correlated with RMC (Supplementary Figure 9). All sequence-driven models that we tested, while highly significant, are only able to explain a minor fraction of the total variance in RMC.

Mechanisms that operate on the RNAs produced by the reporters may play a role in the observed strand asymmetry. In k-mers significantly correlated with RMC, we find motifs associated with RBPs that could affect mRNA steady-state levels (Supplementary Table 2). SNRPB2, HNRNPH2, and RALY are all members of the heterogeneous nuclear ribonucleoprotein (hnRNP) family whose activities include splicing and stabilization of pre-mRNA (Geuens et al. 2016). TIA1 regulates stress granule formation and mRNA aggregation (Rayman and Kandel 2017). CPEB2 and CPEB4 both play roles in poly(A) tail generation resulting in mRNA stabilization (Fernández-Miranda and Méndez 2012). Consistent with this, the A-tail portion of the Alu sequence confers higher RMC (Figure 4). However, none of these mechanisms explain the van Arensbergen et al. results since the test element is not transcribed in that reporter.

We found TF motifs associated with high impact k-mers (Supplementary Table 2). However, the handful of TF motifs found offer only an incomplete view of the potential role of transcriptional regulators in explaining MPRA strand asymmetry. We believe it is likely that both mechanisms that depend on the test element as a template for transcription, and those that depend on its ability to recruit and activate transcriptional complexes are involved. The strong strand asymmetry seen from Alu sequence that contains both RNA stability and RNA polymerase promoter sequences is consistent with this hypothesis. Synthetic mutagenesis of particularly strand asymmetric sequences could allow for the discovery of further sequence content driving the effect.

We have shown the prevalence of strand asymmetric signals from MPRAs in a variety of configurations and contexts. The effect stems from test element sequence, independent of native genomic features due to the nature of the reporters’ construction. We believe that processing of raw MPRA data should be done in strand-aware fashion. Such analyses may yield new biological insights in some cases. Even in cases where only absolute reporter signal from a given sequence is desired, evaluating the maximum value from either strand could reduce false negatives for elements with large strand asymmetries.

Our results make clear that MPRA data detects a biological phenomenon that, while often subtle for a given sequence, is highly reproducible and pervasive across human genomes. The vast multiplexing inherent in MPRA technology has enabled the robust measurements required to find these effects. While further characterization is required, the fact that strand asymmetry is driven by primary sequence and correlates with gene body and Alu element orientation strongly suggests that, whatever the underlying mechanisms are, they are factors that impact gene and genome evolution.

## Methods

### Targeted BAC-derived STARR-seq Assays

We constructed STARR-seq libraries from 14 BACs spanning the HTT locus (Supplementary Table 1). We grew each BAC separately in E. coli, and purified BAC DNA separately according to standard BAC preparation protocols. We sheared each BAC DNA (5 µg) separately using a Biorupter Pico (Diagenode) to 100-500 bp size, then ran on 1% agarose gel, manually selected 250-350 bp size, purified by Qiagen gel extraction, and eluted. Each BAC fragment library separately underwent end repair, dA addition, and paired end adapter ligation (Illumina). Each BAC fragment library separately served as template for PCR with primers FragF (5’-TAGAGCATGCACCGGACACTCTTTCCCTACACGACGCTCTTCCGATCT-3’) and FragR (5’-GGCCGAATTCGTCGACGGTCTCGGCATTCCTGCTGAACCGCTCTTCCGATCT-3’), for 7 or 9 cycles, enough to generate a visible band at target size (∼400 bp). We purified the PCR amplicon libraries by Ampure SPRI beads (Beckman Coulter) and cloned by In-Fusion Cloning (Takara) into the pSTARR-seq_human (Addgene #71509) backbone digested with AgeI and SalI (NEB) according to manufacturers’ protocols. We transformed assembled plasmids by electroporation (Bio-Rad MicroPulser) into MegaX DH10B cells (Thermo Fisher) in four electroporations for each BAC. After recovery, we combined the four cultures for each BAC and grew overnight in 500 mL LB broth. We purified plasmids by the Qiagen Plasmid Maxi Kit. We then pooled plasmid libraries representing each BAC by size of BAC and DNA concentration for equal representation across the locus.

We grew and maintained all cell lines according to ATCC guidelines. For each technical replicate, we transfected 40 million cells 133 ug of reporter plasmid pool. We performed three replicates per cell type. We used Fugene (Promega) as the transfection reagent for A549, BE(2)-C, and HepG2 cells. For K562 cells, we used Lipofectamine PLUS (Invitrogen). After 48 hours, we washed the cells with PBS, and lysed using RLT buffer (Qiagen). We extracted RNA from the lysate using the Total RNA Purification Kit (Norgen), employing four spin columns per replicate and using a lysate volume equivalent to 2 million cells per column. DNA was prepped in a similar manner using the DNEasy Kit (Qiagen). We purified mRNA from the total RNA preps using the DynaBeads poly-A Selection Kit (Invitrogen). We removed contaminating DNA from the mRNA preps using Turbo DNAse I (Ambion). We carried out targeted reverse transcription of reporter RNA using a primer specific to the reporter sequence (5’-CAAACTCATCAATGTATCTTATCATG-3’). Following RNAse treatment, we performed junction PCR for 15 cycles targeting a splice created in the reporter mRNA with the primers: F 5’-GGGCCAGCTGTTGGGGTG*T*C*C*A*C-3’, and R 5’-CTTATCATGTCTGCTCG*A*A*G*C-3’, asterisks indicating phosphorothioate bonds. We prepared sequencing libraries using PCR with Illumina compatible primers and the junction PCR product as template. We generated DNA libraries in the same manner as RNA except that the DNA entered after the reverse transcription step. We sequenced all libraries on the Illumina Next-Seq platform using paired-end 50 bp reads generating ∼40 million reads per replicate on average.

For the *SORT1* locus, we prepped two BACs (Supplementary Table 1) according to standard BAC protocols. We sheared the BAC DNA using a Covaris ultrasonicator and each underwent end repair, dA addition and ligation to custom adaptors:

Left: Starr-adapt-3A 5’-TTGAATTAGATTGATCTAGAGCATGCACCGG*T-3’

Starr-adapt-3C 5’-CCGGTGCATGCTCTAGATCAATC-3’

Right: Starr-adapt-2A 5’-ATGTCTGCTCGAAGCGGCCGGCCGAATTCG*T-3’

Starr-adapt-2C CGAATTCGGCCGGCCGCTTCGAGC.

We size selected ligated fragment on an agarose gel aiming for 1 kB fragments. After gel extraction we subjected the product to 14 cycles of PCR using Starr-adapt-3A and Starr-adapt-2A as primers. We digested the STARR-Seq ORI vector (Addgene 99296) with AgeI and SalI restriction enzymes (NEB), and inserted fragments via NEBuilder HiFi assembly (NEB). For each BAC library, we transformed into NEB #3020 electrocompetent cells and prepped the entire transformation with Chargeswitch Midi Kit (Invitrogen).

We transfected HepGs cells with each BAC-derive library using Lipofectamine (Invitrogen). After 48 hours, we harvested RNA and DNA using the Qiagen Allprep kit (Qiagen). We purified mRNA using the mRNA Mini Kit (Oligotex). We removed contaminating DNA from the mRNA preps using Turbo DNAse I (Ambion). For RNA, we carried out reverse transcription adding a UMI with: P7-StarrBAC-umi-r 5’-CAAGCAGAAGACGGCATACGAGATNNNNNNNNNNCAAACTCATCAATGTATCTTATCATG-3’. This primer also served as the revers primer for PCR of both the cDNA and prepped DNA. The forward primer was: P5-StarrBAC-i# 5’-AATGATACGGCGACCACCGAGATCTACAC##########TGTTGAATTAGATTGATCTAG-3’ where “#” indicates an indexing sequence. We carried out the first round of PCR for 3 cycles and purified the reactions with Ampure XP beads (Beckman-Coulter). We then carried out a second round of PCR with primers targeting the P5 and P7 sequences only: P5 5’-AATGATACGGCGACCACCGAGATCTACA-3’ P7 5’-CAAGCAGAAGACGGCATACGAGAT-3’ for 19-20 cycles. We sequenced the libraries on an Illumina Next-seq generating paired-end 100 bp reads using the custom primers:

StarrBAC-R1 5’-TGTTGAATTAGATTGATCTAGAGCATGCACCGGT-3’

StarrBAC-ind1 5’-GAGCAGACATGATAAGATACATTGATGAGTTTG-3’

StarrBAC-ind2 5’-ACCGGTGCATGCTCTAGATCAATCTAATTCAACA-3’

StarrBAC-R2 5’TCATGTCTGCTCGAAGCGGCCGGCCGAATTCGT-3’

### Sequencing data processing and MPRA signal calculation

We acquired raw FASTQ files from SRA from Johnson et al. (Johnson et al. 2018) (SRP144640), using sratoolkit (v2.9.6-1). We aligned raw paired-end Illumina reads to either genome (hg38) subsets corresponding to the regions of BAC coverage for BAC-derived libraries, or to the whole genome using bowtie2 (v 2.2.5) (Langmead and Salzberg 2012). For BAC derived libraries, we also included the E. coli genome (K-12 MG1655) in the reference to filter out E. coli genomic DNA contaminants and assess BAC prep purity. Using the alignment positions and flag sum in the aligned BAMs, we constructed BED files of the sequenced fragment including its orientation to reference using samtools v 1.8 (Li et al. 2009) and a custom PERL script. We have included the scripts that take FASTQs to fragment BED files in Supplementary Files. We refer to fragments aligning to the reference as “Reference” and those to the reference reverse complement as “Complement”.

We created a set of 290 bp non-overlapping bins spanning the autosome using the R (v3.6.1) package GenomicRanges v1.36 (Lawrence et al. 2013). We picked 290 bp because it was the median fragment size of the BAC-derived libraries. For BAC-derived libraries, we reduced the bins to only those that overlapped the BAC-covered regions to facilitate computation. We converted the sequencing data fragment BED files to GenomicRanges objects using the R package rtracklayer v1.44.3 (Lawrence et al. 2009). We found overlaps of Reference and Complement fragments separately for each bin, generating bin counts from each strand. We supply the scripts for generating bin counts in Supplementary Files.

Each experiment had DNA counts corresponding to reporter input levels and RNA counts corresponding to transcripts derived from the reporters. We normalized the raw counts of each pair of DNA and RNA versus strand by dividing by the sum of each count type across all bins. We then calculated the reporter activity for each strand by dividing the normalized RNA counts by the normalized DNA counts. To measure strand asymmetry, we subtracted the Complement strand signal from the Reference strand signal to yield Reference minus Complement (RMC).

The reporter data from van Arensbergen et al employed barcodes associated by a separate sequencing run with upstream elements (van Arensbergen et al. 2019). Because of the complexities of associating barcodes to test elements in an unfamiliar experimental design we did not perform ourselves, we instead downloaded the stranded reporter signal bigWigs from the metadata in the GEO submission (GSE128325). These bigWigs are also hg19 referenced, so we mapped with the import function in rtracklayer the bigWig values to the autosome bin set that we had lifted from hg38 to hg19 using the UCSC Browser liftover tool (Kent et al. 2002). After mapping, we back converted to hg38 so that this data would be comparable to the other datasets. We calculated RMC for this data as above but noticed outliers of very high absolute RMC. HMMSeg assumes a Gaussian underlying distribution and these outliers interfered with the segmentation calculations. Accordingly, we divided the RMC values by the maximum value of either strand (RMC over Max) and found that the outliers and overall variance between binned values in test groupings (such as Reference gene bodies) were reduced.

### Hierarchical clustering and association with gene bodies

To create heatmaps of data hierarchically clustered by similarity, we calculated the Spearman correlation coefficient, rho, between each sample within an experiment. We then calculated the Euclidean distance between each sample and clustered using the R functions dist and hclust. We created heatmaps with the indicated saturation color ranges.

For the Johnson et al. data, we observed agreement in strand asymmetry across dexamethasone treatment duration. In order to have more accurate genome-wide data, we summed the sequencing bin counts across all dexamethasone treatment durations to a single RMC measurement for the dataset. We filtered out bins with less than 57 summed DNA counts, which we calculated would yield 10 RNA counts from each strand for a neutral test element, on average. For the van Arensbergen et al. data, we noticed similar agreement across donor genomic DNA. For this data, we took the median signal across the donors and calculated a single RMC over Max for each cell type.

To compare these data to gene bodies, we constructed a GenomicRanges object for gene models obtained from GTEx v8 without limiting to protein coding or any other filter. All regions of the autosome without an annotated gene model we labeled “intergenic”. We labeled regions creating sense transcripts matching reference “Reference”. Those with sense transcripts matching the reverse complement of reference we labeled “Complement”. Wherever there were annotated gene models on both strands, we labeled them “Opposite overlapping”. In this way we divided the entire autosome into four categories. To evaluate the correlation of RMC or RMC over Max with these gene body categories, we calculated the overlap of each autosome bin to each category. Using the category as the independent variable and the RMC or RMC over Max as the dependent variable, we performed linear regression using the R function lm. Boxplots were made in a similar fashion.

To create segmentations of the RMC or RMC over Max values, we employed a Hidden Markov Model approach via the HMMSeg software package (Day et al. 2007). For all data we used a two state model. For the whole genome datasets, the emission means and variances were calculated from the means and variances of the data in Complement and Reference genes, respectively. Because the BAC-derived datasets encompassed smaller regions containing a small number of gene models, we used the emission means and variances of the Johnson et al. data in the segmentation models of these. For all data sets, we tested a range of transition probabilities from 0.05 to 0.5. We present data from a representative subset of these. To evaluate the similarity of the calculated segmentations to gene body classes, we employed a conditional entropy approach (Haiminen et al. 2007). We calculated the conditional entropy (H) of the HMM-seq segmentation (P) given a stranded gene body class, for example Reference genes (Q), based on the lemma H(P|Q) = H(U) – H(Q) provided in Haiminen et al. where U is the union of all segment borders in both P and Q. We then isentroprically shuffled P (maintaining the width and number of segments but shuffling start positions) 1,000 times and calculated H(P|Q) of each shuffle. We evaluated significance by comparing the actual value of H(P|Q) to the distribution of values from shuffles. We employed the process separately to Reference and Complement genes, considering both p-values in our evaluation of significance.

### Calculation of Alu sequence effect

We downloaded the complete BED file of RepeatMasker tracks from the UCSC Browser (Kent et al. 2002). Using a custom Perl script, we processed this file to pull out Alu positions creating a BED file of every Alu base with genomic position, position within the Alu consensus sequence, strand, and divergence (in milliDiv units). To this Alu reference, we then counted overlaps of the fragment BED files from Johnson et al. data in the same process used for the autosome bins. Many (∼1 million) genomic positions map to each Alu consensus base. We split each consensus base into three blocks of divergence by milliDiv thresholds of <100, 100-200, and >200. Then we summed stranded counts from genomic positions to their Alu consensus position / milliDiv group. In this case we kept the dexamethasone treatment sets separate in order to estimate variance and because the subsequent collapsing to consensus sequence yields sufficiently numerous counts. From these collapsed counts, we calculated median RMC at each Alu consensus position / milliDiv group set as well as the standard deviation across dexamethasone treatment durations.

### Sequence-based modeling of MPRA asymmetry

To build a linear model relating sequence content to RMC, we first counted the frequency of all monomers, dimers, and octamers in each bin across the genome, allowing k-mer overlaps. For each k-mer, we then performed a genome-wide Spearman’s correlation between RMC and k-mer count on three genome bin sets: All bins, the combination of top 1% and bottom 1% RMC bins, and the middle 80% RMC bins. To avoid overfitting and make a linear model computationally tractable, we reduced the set of octamers to fewer than 1000 octamers by selecting the maximum p-value between the three correlations for each octamer, and using a p-value cutoff of 1E-230. This reduced the list to 936 octamers. We then used the counts for each of these 936 octamers, all dimers, and all monomers (for a total of 956 k-mers) as predictor variables in a linear model of RMC ∼ kmer Counts. The model was trained on all data from chromosome 1, and tested on all data from chromosome 2. Additionally, we created a model for each individual k-mer using that k-mer’s count as the sole independent variable to determine individual r^2^ values. For comparison to RBP and TF motifs within the CISBP database (Weirauch et al. 2014), we identified all octamers which were found significant in the linear model. We saved these octamers as a FASTA file, and used the FIMO function (Grant et al. 2011) of the meme suite (version 5.1.0), using default parameters.

We ran Xpresso using a pre-trained convolutional neural network model intended to predict median gene expression levels across cell types (https://xpresso.gs.washington.edu/) (Agarwal and Shendure 2020). We computed the predicted RMC as the difference between the predicted value of Xpresso run on the Reference versus Complement strand, centered upon the same intervals used to calculate RMC from MPRA data.

## Data Access

All raw and processed sequencing data generated in this study have been submitted to the NCBI Gene Expression Omnibus (GEO; https://www.ncbi.nlm.nih.gov/geo/) under accession number GSE156857.

## Acknowledgements

We would like to thank the Tim Reddy lab at Duke University and the Bas van Steensel lab at the Netherlands Cancer Institute for their generation of high-quality MPRA datasets. This work was supported by the CHDI Foundation (to R.M.), the Leo Fund at HudsonAlpha, and National Institutes of Health (NIH) grants 1UM1HG009408 and 1R01HG009136 (to J.S.). Jay Shendure is an investigator of the Howard Hughes Medical Institute.

## References

Agarwal V, Shendure J. 2020. Predicting mRNA Abundance Directly from Genomic Sequence Using Deep Convolutional Neural Networks. Cell Rep 31: 107663. https://doi.org/10.1016/j.celrep.2020.107663.

Almada AE, Wu X, Kriz AJ, Burge CB, Sharp PA. 2013. Promoter directionality is controlled by U1 snRNP and polyadenylation signals. Nature 499: 360–363.

Andersson R, Chen Y, Core L, Lis JT, Sandelin A, Jensen TH. 2015. Human Gene Promoters Are Intrinsically Bidirectional. Mol Cell 60: 346–347. http://dx.doi.org/10.1016/j.molcel.2015.10.015.

Andersson R, Gebhard C, Miguel-Escalada I, Hoof I, Bornholdt J, Boyd M, Chen Y, Zhao X, Schmidl C, Suzuki T, et al. 2014. An atlas of active enhancers across human cell types and tissues. Nature 507: 455–461.

Arnold CD, Gerlach D, Stelzer C, Boryń ŁM, Rath M, Stark A. 2013. Genome-wide quantitative enhancer activity maps identified by STARR-seq. Science 339: 1074–1077. http://www.ncbi.nlm.nih.gov/pubmed/23328393.

Ashe HL, Monks J, Wijgerde M, Fraser P, Proudfoot NJ. 1997. Intergenic transcription and transinduction of the human β-globin locus. Genes Dev 11: 2494–2509.

Barakat TS, Halbritter F, Zhang M, Rendeiro AF, Perenthaler E, Bock C, Chambers I. 2018. Functional Dissection of the Enhancer Repertoire in Human Embryonic Stem Cells. Cell Stem Cell 23: 276–288.e8.

Bell AC, West AG, Felsenfeld G. 1999. The protein CTCF is required for the enhancer blocking activity of vertebrate insulators. Cell 98: 387–396.

Dao LTM, Galindo-Albarrán AO, Castro-Mondragon JA, Andrieu-Soler C, Medina-Rivera A, Souaid C, Charbonnier G, Griffon A, Vanhille L, Stephen T, et al. 2017. Genome-wide characterization of mammalian promoters with distal enhancer functions. Nat Genet 49: 1073–1081.

Day N, Hemmaplardh A, Thurman RE, Stamatoyannopoulos JA, Noble WS. 2007. Unsupervised segmentation of continuous genomic data. Bioinformatics 23: 1424–6. http://www.ncbi.nlm.nih.gov/pubmed/17384021 (Accessed July 25, 2017).

Deininger P. 2011. Alu elements: Know the SINEs. Genome Biol 12: 1–12.

Dekker J, Rippe K, Dekker M, Kleckner N. 2002. Capturing chromosome conformation. Science (80-) 295: 1306–1311.

Dunham I, Kundaje A, Aldred SF, Collins PJ, Davis CA, Doyle F, Epstein CB, Frietze S, Harrow J, Kaul R, et al. 2012. An integrated encyclopedia of DNA elements in the human genome. Nature 489: 57–74.

Duttke SHC, Lacadie SA, Ibrahim MM, Glass CK, Corcoran DL, Benner C, Heinz S, Kadonaga JT, Ohler U. 2015. Human promoters are intrinsically directional. Mol Cell 57: 674–684. http://dx.doi.org/10.1016/j.molcel.2014.12.029.

Fernández-Miranda G, Méndez R. 2012. The CPEB-family of proteins, translational control in senescence and cancer. Ageing Res Rev 11: 460–472. http://dx.doi.org/10.1016/j.arr.2012.03.004.

Geuens T, Bouhy D, Timmerman V. 2016. The hnRNP family: insights into their role in health and disease. Hum Genet 135: 851–867.

Gordon MG, Inoue F, Martin B, Schubach M, Agarwal V, Whalen S, Feng S, Zhao J, Ashuach T, Ziffra R, et al. 2020. lentiMPRA and MPRAflow for high-throughput functional characterization of gene regulatory elements. Nat Protoc 15. http://dx.doi.org/10.1038/s41596-020-0333-5.

Grant CE, Bailey TL, Noble WS. 2011. FIMO: Scanning for occurrences of a given motif. Bioinformatics 27: 1017–1018.

Green P, Ewing B, Miller W, Thomas PJ, Thomas J, Touchman J, Blakesley R, Bouffard G, Beckstrom-Sternberg S, McDowell J, et al. 2003. Transcription-associated mutational asymmetry in mammalian evolution. Nat Genet 33: 514–517.

Haiminen N, Mannila H, Terzi E. 2007. Comparing segmentations by applying randomization techniques. BMC Bioinformatics 8: 1–8.

Johnson DS, Mortazavi A, Myers RM, Wold B. 2007. Genome-Wide Mapping of in Vivo Protein-DNA Interactions. Science (80-) 316: 1497–1502.

Johnson GD, Barrera A, McDowell IC, D’Ippolito AM, Majoros WH, Vockley CM, Wang X, Allen AS, Reddy TE. 2018. Human genome-wide measurement of drug-responsive regulatory activity. Nat Commun 9: 1–9. http://dx.doi.org/10.1038/s41467-018-07607-x.

Jurka J, Kapitonov V V, Klonowski P, Walichiewicz J, Smit AF. 1996. Identification of new medium reiteration frequency repeats in the genomes of Primates, Rodentia and Lagomorpha. Genetica 98: 235–247.

Kent WJ, Sugnet CW, Furey TS, Roskin KM, Pringle TH, Zahler AM, Haussler a. D. 2002. The Human Genome Browser at UCSC. Genome Res 12: 996–1006.

Klein J, Agarwal V, Inoue F, Keith A, Martin B, Kircher M, Ahituv N, Shendure J. 2019. A systematic evaluation of the design, orientation, and sequence context dependencies of massively parallel reporter assays. bioRxiv 576405.

Kwasnieski JC, Mogno I, Myers CA, Corbo JC, Cohen BA. 2012. Complex effects of nucleotide variants in a mammalian cis-regulatory element. Proc Natl Acad Sci U S A 109: 19498–19503.

Langmead B, Salzberg SL. 2012. Fast gapped-read alignment with Bowtie 2. Nat Methods 9: 357–359.

Lawrence M, Gentleman R, Carey V. 2009. rtracklayer: An R package for interfacing with genome browsers. Bioinformatics 25: 1841–1842.

Lawrence M, Huber W, Pagès H, Aboyoun P, Carlson M, Gentleman R, Morgan MT, Carey VJ. 2013. Software for Computing and Annotating Genomic Ranges. PLoS Comput Biol 9: 1–10.

Li H, Handsaker B, Wysoker A, Fennell T, Ruan J, Homer N, Marth G, Abecasis G, Durbin R. 2009. The Sequence Alignment/Map format and SAMtools. Bioinformatics 25: 2078–2079.

Liu Y, Yu S, Dhiman VK, Brunetti T, Eckart H, White KP. 2017. Functional assessment of human enhancer activities using whole-genome STARR-sequencing. Genome Biol 18: 1–13.

Mikhaylichenko O, Bondarenko V, Harnett D, Schor IE, Males M, Viales RR, Furlong EEM. 2018. The degree of enhancer or promoter activity is reflected by the levels and directionality of eRNA transcription. Genes Dev 32: 42–57.

Mills RE, Bennett EA, Iskow RC, Devine SE. 2007. Which transposable elements are active in the human genome? Trends Genet 23: 183–191.

Moore JE, Purcaro MJ, Pratt HE, Epstein CB, Shoresh N, Adrian J, Kawli T, Davis CA, Dobin A, Kaul R, et al. 2020. Expanded encyclopaedias of DNA elements in the human and mouse genomes. Nature 583: 699–710. http://www.nature.com/articles/s41586-020-2493-4.

Muerdter F, Boryń ŁM, Arnold CD. 2015. STARR-seq – Principles and applications. Genomics 106: 145–150.

Patwardhan RP, Lee C, Litvin O, Young DL, Pe’Er D, Shendure J. 2009. High-resolution analysis of DNA regulatory elements by synthetic saturation mutagenesis. Nat Biotechnol 27: 1173–1175.

Plank JL, Dean A. 2014. Enhancer function: Mechanistic and genome-wide insights come together. Mol Cell 55: 5–14. http://dx.doi.org/10.1016/j.molcel.2014.06.015.

Ramaker RC, Hardigan AA, Goh ST, Partridge EC, Wold B, Cooper SJ, Myers RM. 2020. Dissecting the regulatory activity and sequence content of loci with exceptional numbers of transcription factor associations. Genome Res 30: 939–950.

Rayman JB, Kandel ER. 2017. TIA-1 Is a Functional Prion-Like Protein. 1–14.

Schoenfelder S, Fraser P. 2019. Long-range enhancer–promoter contacts in gene expression control. Nat Rev Genet 20: 437–455. http://dx.doi.org/10.1038/s41576-019-0128-0.

Schöne S, Bothe M, Einfeldt E, Borschiwer M, Benner P, Vingron M, Thomas-Chollier M, Meijsing SH. 2018. Synthetic STARR-seq reveals how DNA shape and sequence modulate transcriptional output and noise. PLoS Genet 14: 1–24.

Sherwood RI, Hashimoto T, O’Donnell CW, Lewis S, Barkal AA, Van Hoff JP, Karun V, Jaakkola T, Gifford DK. 2014. Discovery of directional and nondirectional pioneer transcription factors by modeling DNase profile magnitude and shape. Nat Biotechnol 32: 171–178. http://www.nature.com/articles/nbt.2798 (Accessed July 12, 2018).

Sun J, He N, Niu L, Huang Y, Shen W, Zhang Y, Li L, Hou C. 2019. Global Quantitative Mapping of Enhancers in Rice by STARR-seq. Genomics, Proteomics Bioinforma 17: 140–153.

van Arensbergen J, Pagie L, FitzPatrick VD, de Haas M, Baltissen MP, Comoglio F, van der Weide RH, Teunissen H, Võsa U, Franke L, et al. 2019. High-throughput identification of human SNPs affecting regulatory element activity. Nat Genet 51. http://dx.doi.org/10.1038/s41588-019-0455-2.

Visel A, Blow MJ, Li Z, Zhang T, Akiyama JA, Holt A, Plajzer-Frick I, Shoukry M, Wright C, Chen F, et al. 2009a. ChIP-seq accurately predicts tissue-specific activity of enhancers. Nature 457: 854–858.

Visel A, Rubin EM, Pennacchio LA. 2009b. Genomic views of distant-acting enhancers. Nature 461: 199–205.

Wang X, He L, Goggin SM, Saadat A, Wang L, Sinnott-Armstrong N, Claussnitzer M, Kellis M. 2018. High-resolution genome-wide functional dissection of transcriptional regulatory regions and nucleotides in human. Nat Commun 9. http://dx.doi.org/10.1038/s41467-018-07746-1.

Weirauch MT, Yang A, Albu M, Cote AG, Montenegro-Montero A, Drewe P, Najafabadi HS, Lambert SA, Mann I, Cook K, et al. 2014. Determination and inference of eukaryotic transcription factor sequence specificity. Cell 158: 1431–1443.

